# Metabolic design in a model of extreme mammalian metabolism, the North American least shrew (*Cryptotis parva*)

**DOI:** 10.1101/2021.05.28.446190

**Authors:** Dillon J. Chung, Grey P. Madison, Angel M. Aponte, Komudi Singh, Yuesheng Li, Mehdi Pirooznia, Christopher K. E. Bleck, Nissar A. Darmani, Robert S. Balaban

## Abstract

Mitochondrial adaptations are fundamental to differentiated function and energetic homeostasis in mammalian cells. But the mechanisms that underlie these relationships remain poorly understood. Here, we investigated organ-specific mitochondrial morphology, connectivity and protein composition in a model of extreme mammalian metabolism, the Least shrew (*Cryptotis parva*). This was achieved through a combination of high-resolution 3D focused-ion-beam EM imaging and tandem-mass-tag MS proteomics. We demonstrate that liver and kidney mitochondrial content are equivalent to the heart permitting assessment of mitochondrial adaptations in different organs with similar metabolic demand. Muscle mitochondrial networks (cardiac and skeletal) are extensive, with a high incidence of nanotunnels – which collectively support the metabolism of large muscle cells. Mitochondrial networks were not detected in the liver and kidney as individual mitochondria are localized with sites of ATP consumption. This configuration is not observed in striated muscle, likely due to a homogenous ATPase distribution and the structural requirements of contraction. These results demonstrate distinct, fundamental mitochondrial structural adaptations for similar metabolic demand that are dependent on the topology of energy utilization process in a mammalian model of extreme metabolism.

## Introduction

Organisms that maintain homeostasis at physiological extremes are powerful models in which to investigate the mechanisms used to regulate metabolism as the evolutionary responses that they exhibit potentially reveal processes most critical to survival (1–3). One way to identify species with extreme metabolic rates is to exploit the phenomenon of allometric mass-scaling of metabolism (i.e., the hypometric relationship between mass-specific metabolic rate and organism size) (4,5). These mass-scaling relationships are greatest under basal metabolic conditions (4), which suggests that organs that are the greatest contributors to this state (e.g., liver, kidney) are most likely to exhibit metabolic modifications that underpin extreme basal metabolism in small organisms. The North American least shrew (*Cryptotis parva*) is an ideal mammalian model in which to study these adaptations as these small animals (body mass = 4-6 g) (6) are positioned at the far end of the allometric scale – achieving among the highest mass-specific metabolic rates (MO_2 resting_ = 4 mL O_2_ min^-1^ g^-1^) (7) and heart rates (HR_resting_ = 777 min^-1^) (8). In addition, *C. parva* derives from a different Order (*Eulipotyphla*) than the extensively studied mouse (*Rodentia*) potentially providing a different evolutionary path for the design of energy metabolism. Despite this impressive physiology, the mechanisms that underpin and sustain the extreme metabolism of small mammals – such as *C. parva* – remain poorly understood.

Mitochondrial performance is integral to the maintenance of mammalian energetic homeostasis. Thus, the high basal metabolic rates exhibited by small organisms should be associated with mitochondrial characteristics that enhance function to meet cellular demand – particularly in tissues associated with basal metabolism. Mitochondrial capacity can be manipulated through mechanisms that include increased mitochondrial content, altered mitochondrial enzyme composition/post-translational modifications, and modifications to mitochondrial structure and interactions. There is good evidence that electron transport system (ETS) enzyme content increases in a tissue-specific manner with decreasing organism size (9–11). These scaling effects are largest in the liver and smaller effects are observed in organs associated with maximum metabolism such as the heart (11) – a tissue-specific pattern that is consistent with the scaling of tissue-level respiration (12–14). However, it remains unclear whether the greater ETS enzyme content observed in small mammals is a proteome-wide phenomenon and what influence tissue-specific demands impose on these scaling responses. Tissue-specific scaling effects are also observed in measurements of mammalian mitochondrial volume density (15,16). It has recently been shown that murine skeletal muscle and cardiac mitochondria form networks that distribute the mitochondrial proton motive force over intracellular distances which potentially aids in the maintenance of cellular energetic homeostasis (17). But the relevance of mitochondrial connectivity as a mechanism that maintains the extreme metabolic demand of smaller organisms remains unknown and it is unclear whether mitochondrial networking is present or functionally important in non-contractile tissues.

Here, we compared mitochondrial content, morphology, network architecture and protein composition among tissues that represent the largest contributors to basal and maximum metabolism (i.e., heart, skeletal muscle, liver and kidney) in captive-bred *C. parva*. This tissue-specific comparison was employed as mitochondrial performance varies according to the demands of each tissue – potentially resulting in functional trade-offs that become apparent in the mitochondria of small organisms. We utilized focused ion beam-SEM imaging and developed segmentation workflows to achieve nanometer-scale 3D quantification of mitochondrial interactions. In addition, we employed a tandem-mass-tag proteomics approach to assess protein programming among these organs. This proteomics screen incorporated a *de novo* sequenced and annotated *C. parva* draft genome as part of its database.

Through this work, we demonstrate that mitochondrial content and protein programming in the liver and kidney approach levels observed in the metabolically active heart of *C. parva* – a configuration that likely supports the extreme metabolic rates of small mammals. We identify mitochondrial-ER interactions and ion pumping as processes that are sustained by this high mitochondrial content and are thus contributors to the allometric scaling of mammalian metabolic rates. Mitochondrial networks are limited to striated muscle in *C. parva* and are oriented in ways that are consistent with those observed in larger mammals. But these networks also exhibit increased connectivity at the nm scale which may reflect the high metabolic requirements of this species. Altogether, we determine that mitochondrial network formation is a consequence of the contractile requirements of the musculature and not mitochondrial content or tissue metabolic demand – a finding that improves our fundamental understanding of the mitochondrial properties that are altered to maintain mammalian energetic homeostasis.

## Results

### Mitochondrial volume density is greater in *C. parva* than in larger mammals

Alterations to mitochondrial content are a simple mechanism through which organisms can meet diverse cellular energetic requirements. Using nanometer scale FIB-SEM imaging (voxel size = 10 x 10 x 10 nm) and high-volume AI segmentation (Fig. 1, Movie S1) we compared mitochondrial volume density among the skeletal muscle, heart, liver and kidney as these organs have vastly different functions and are on the extreme ends of tissues that contribute to maximum and basal metabolic rates (4,16).

**Figure. 1.**
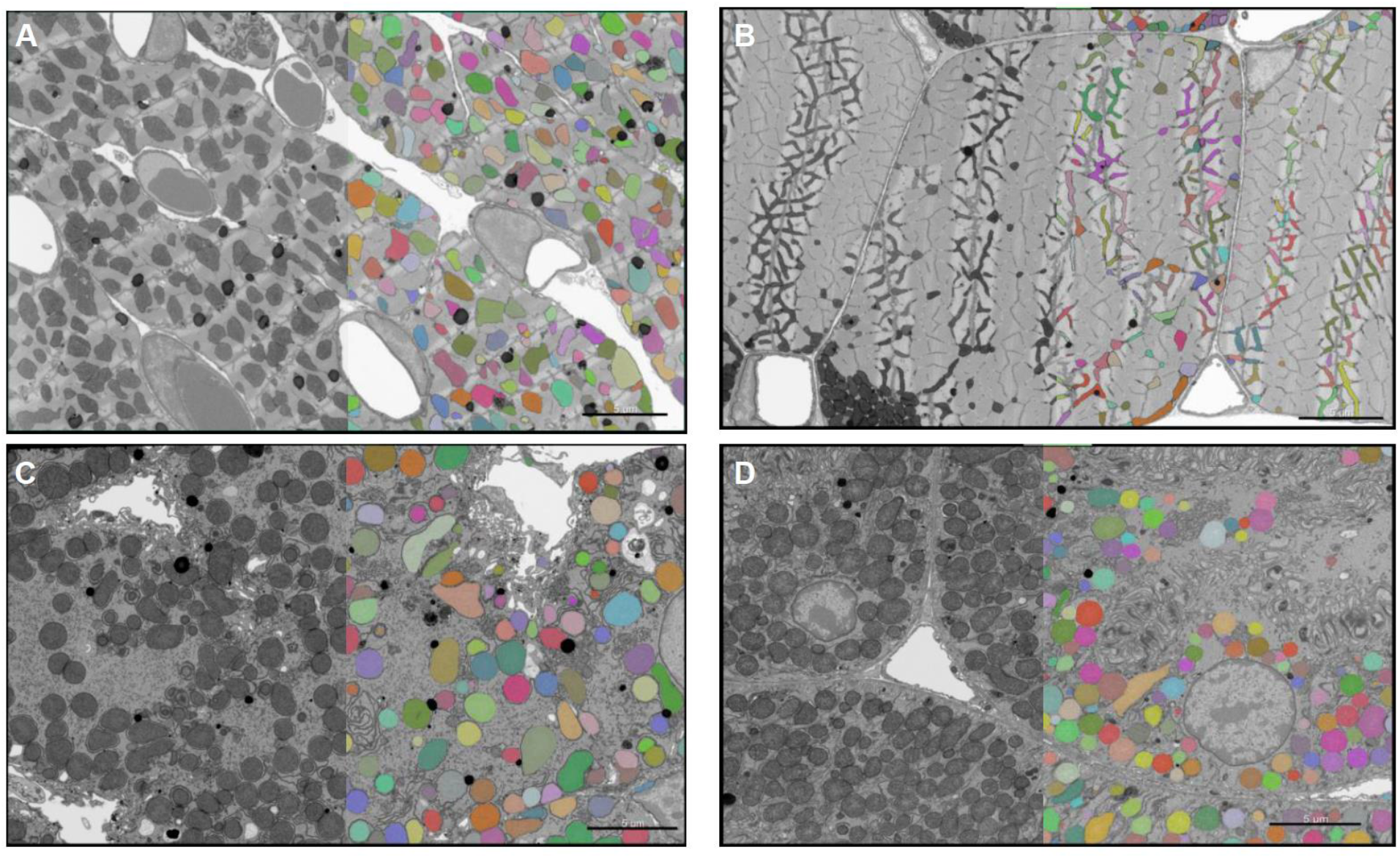
Representative segmentation of mitochondria from four tissues in *Cryptotis parva*. Data are raw FIB-SEM images (greyscale) with an overlay of mitochondrial segmentation (false color) from heart (**A**), skeletal muscle (**B**), liver (**C**), and kidney (**D**). Distinct colors indicate individually segmented mitochondria. Scale bar: 5 μm.

Cardiac mitochondrial volume density is relatively insensitive to changes in body size and our data are consistent with this phenomenon (9,18,19). This near-isometric relationship between cardiac mitochondrial volume density and organism size indicates that cardiac design is optimized for energy conversion and myofilament function across mammals (11). In contrast to the mass scaling of heart mitochondria – liver and kidney mitochondrial volume density and enzymatic content increase substantially with decreasing organism size and this is supported by comparisons of our data with those from larger mammals (9,11) and the similar mitochondrial volume density between these tissues (liver: 36.1 ± 1.07%, kidney: 40.2 ± 1.82%) and the heart (33.2 ± 0.72%) (Fig. 2M). The presence of high mitochondrial volume density in the liver and kidney provides evidence that processes in these organs drive the extreme basal metabolism of *C. parva* and small mammals more generally.

**Figure 2.**
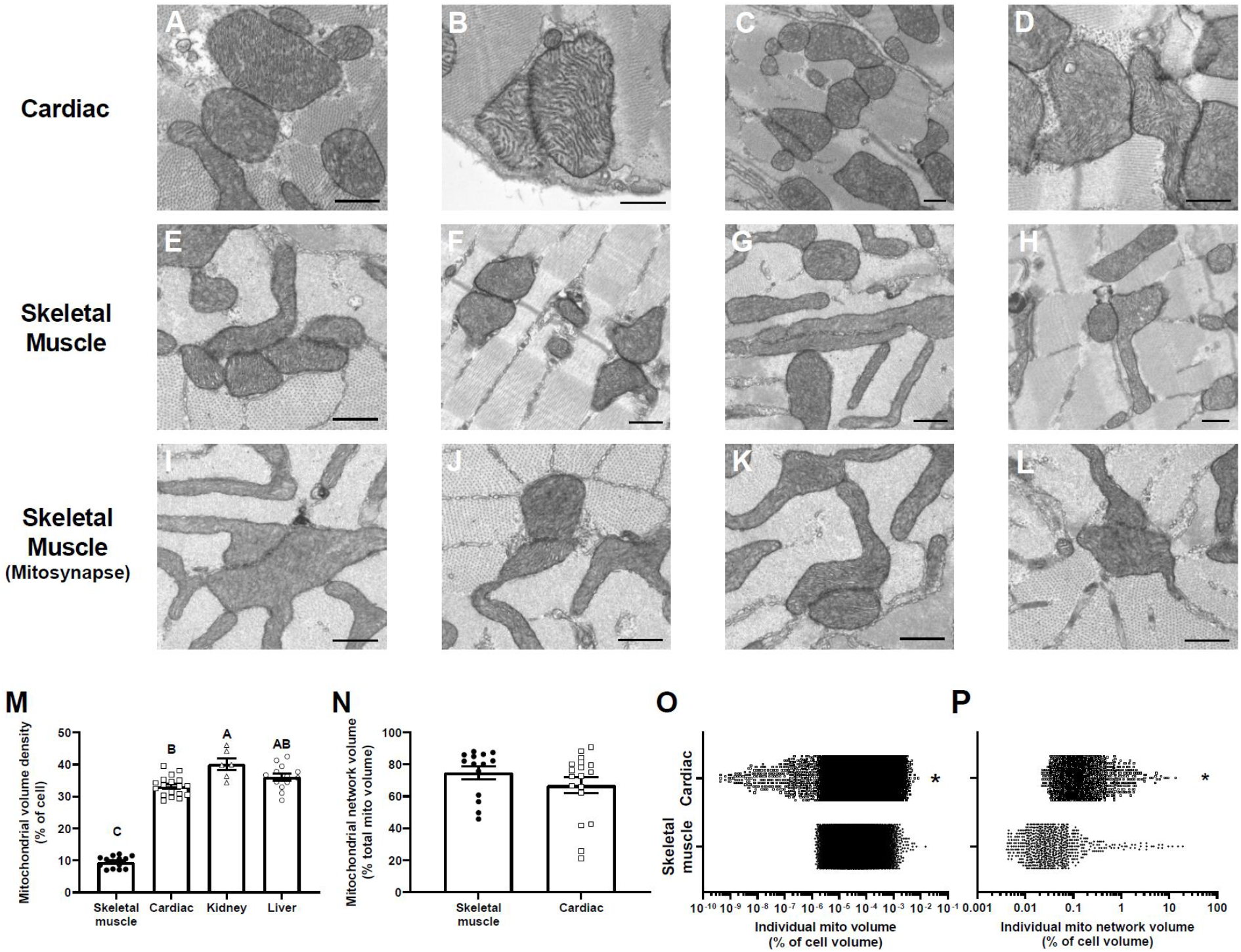
*Cryptotis parva* exhibits extreme mitochondrial content and high mitochondrial connectivity. Intermitochondrial junctions (IMJs) are prevalent in the heart (**A-D**) and skeletal muscle (**E-H**). Some skeletal muscle intermyofibrillar IMJs form unique structures with high outer mitochondrial membrane surface area – termed mitosynapses (**I-L**). Data are TEM images; scale bar: 500 nm. **M**) Mitochondrial volume density. **N**) Mean volume density of IMJ-linked mitochondrial networks. Note the increase in volume density between individual mitochondria (**O**) and IMJ-linked mitochondrial networks (**P**). Different letters indicate a significant difference among tissues (ANOVA or Dunn’s multiple comparison, p < 0.05). Asterisks indicate a significant difference between muscle type (Tukey or Mann-Whitney, p < 0.05). Summary data are mean ± SEM. Skeletal muscle, *n* = 15646 mitochondria, 16 datasets, three animals; cardiac muscle, *n* = 15893 mitochondria, 18 datasets, three animals; kidney, *n* = 16143 mitochondria, six datasets, two animals; liver, *n* = 6939 mitochondria, 13 datasets, three animals. Skeletal muscle networks, *n* = 682 networks, 16 datasets, three animals; cardiac muscle, *n* = 1218 networks, 18 datasets, three animals.

Skeletal muscle mitochondrial volume density (9.48 ± 0.49%) is low compared to other sampled tissues in *C. parva* (Fig. 2M). Like cardiac muscle, skeletal muscle mitochondrial volume density exhibits a near isometric mass-scaling relationship which is similar to the mass-scaling of maximum metabolic rates and is consistent with the contribution of these tissues to this metabolic state (9,20). In contrast to data from *M. musculus* (19), we do not observe considerable variability in mitochondrial volume density between cells with high fiber-parallel mitochondrial (FPM) content – a characteristic often used to identify more ‘aerobic’ cells (19) – and cells with low FPM (‘glycolytic’ cells; Fig. 2M). This suggests that *C. parva* skeletal muscle cells exhibit a more homogenous fiber type – a conclusion that is supported by the indistinguishable myosin chain composition and fiber type pattern between extensor digitorum longus and soleus muscle in Etruscan (*Suncus etruscus*) and European white-tooth shrews (*Crocidura russula*) (21) and the increased oxidative and decreasing glycolytic enzyme activities in gastrocnemius muscle with decreasing size in mammals (22). The similar mitochondrial volume density but contrasting FPM content in these cells could also reflect variation in the localization of energetic demand of cells that participate in chronic (high FPM) versus burst (low FPM) activity.

We note that some skeletal muscle cells exhibit an exceptionally high mitochondrial content (Fig. S1A). These outliers (22.2 and 30.3% mitochondrial volume density) are likely due to increased sampling of paravascular and paranuclear mitochondrial pools as indicated by the high average mitochondrial sphericity in these cells (Fig S1D). It is probable that current quantification methods underestimate skeletal muscle mitochondrial volume density due to a higher likelihood of sampling intermyofibrillar (IMF) regions. The high mitochondrial content coupled with an electrically conductive reticulum hints at the prevalence and functional importance of heterogeneous mitochondrial distributions within skeletal muscle cells that maintain energetic homeostasis with minimal impact on the fiber geometry required for contraction (23).

### Mitochondrial connectivity is high in skeletal and cardiac muscle and is not present in the liver and kidney of *Cryptotis parva*

Mitochondrial reticulation has gained attention as a mechanism that maintains energetic homeostasis across intracellular distances (17,24). We predicted that the extreme sustained metabolic requirements of *C. parva* would result in extensive mitochondrial networking – achieved through a greater incidence of direct mitochondrial contacts (intermitochondrial junctions: IMJ; mitosynapses) and extensions of the mitochondrial matrix (tunneling). We combined 3D segmentation of mitochondria and IMJs – structures that are proposed to conduct mitochondrial proton motive force across adjacent outer mitochondrial membranes – to quantify mitochondrial matrix continuity in skeletal muscle and heart (Fig. 2A – L; Movie S2, S3) (19,25). We demonstrate that *C. parva* maintain mitochondrial interactions in the heart and skeletal muscle at levels greater than those observed in larger organisms (Fig. 2, 3). However, mitochondrial reticulation is not observed in the liver and kidney (Fig. 3E-F; Movie S1C - D) despite the high mitochondrial volume density and predicted resting metabolic rates in these diminutive animals. When combined with tissue-specific differences in mitochondrial volume density, we propose that the presence of a mitochondrial reticulum is not a consequence of the metabolic demand of the tissue or mitochondrial density but is instead a function of spatial constraints imposed by the contractile elements of musculature.

**Figure 3.**
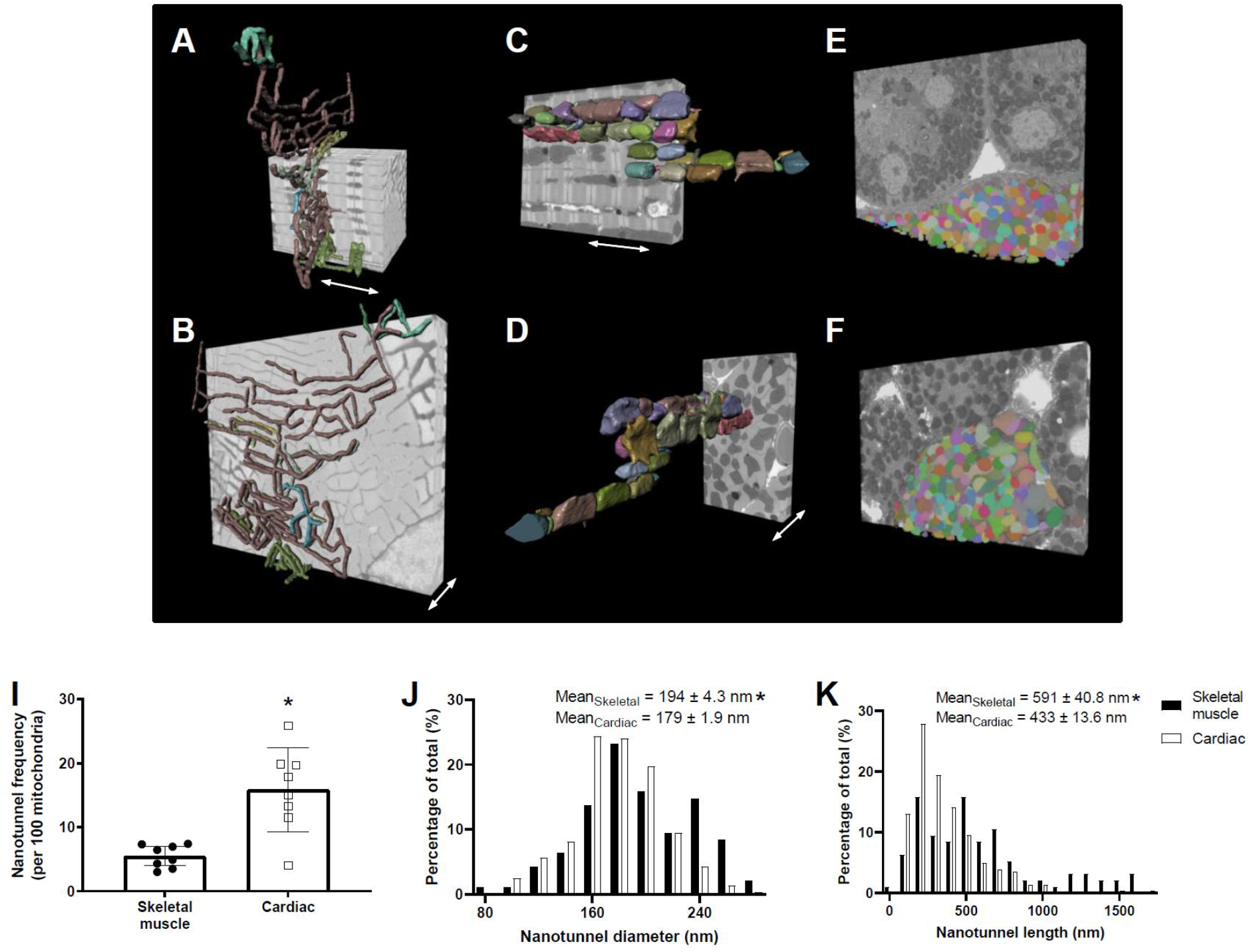
Mitochondrial networks are limited to striated muscle and recruit nanotunnels as part of their structure. 3D renderings of a single mitochondrial network in skeletal muscle (**A-B**) and heart (**C-D**), oriented parallel (A, C) or perpendicular (B, D) to the plane of contraction (double arrow). Mitochondrial networks are not observed in the kidney (**E**) or liver (**F**) and mitochondria are distributed throughout the cytosol in these tissues. Distinct colors indicate individual mitochondria and 3D renderings are partially overlaid on FIB-SEM volumes. **I**) Mitochondrial nanotunnel frequency. **J**) Nanotunnel diameter. **K**) Nanotunnel length. Asterisks indicate a significant difference between tissues (Tukey or Mann-Whitney, p < 0.05). Summary data are mean ± SEM. Skeletal muscle, *n* = 100 nanotunnels, 8 datasets, two animals; cardiac muscle, *n* = 286 nanotunnels, 8 datasets, three animals.

We confirm that striated muscle mitochondrial networks are organized in perpendicular (skeletal muscle) and parallel (cardiac) orientations relative to the plane of contraction as observed in other mammalian systems (17,19) (Fig. 3A – D; Movie S3, S4). In skeletal muscle, the mitochondrial reticulum is primarily linked through matrix continuity, with IMJs connecting I-band mitochondria to peripheral mitochondrial pools (Movie S4B, S5B). These I-band mitochondria differ from those observed in larger mammals (19,26) in that they form flattened tube-shaped structures that may act to increase outer mitochondrial membrane surface area (Movie S3C – D). Many skeletal muscle IMJs form what appear to be specialized structures with both sides of the IMJ exhibiting large contact zones reminiscent of neuronal synapses (Fig. 2I - L). Depending on the region being examined, several of these structures can be observed in a single plane. These structures are consistent with a local increase in surface contact between mitochondria and are termed mitosynapses.

In cardiac muscle, I-band mitochondria are not present – instead, FPM are connected end-to-end via IMJs with lateral nanotunneling that connects FPM pools separated by myofibrils (Fig. 3C – D, Movie S3A - B, S4A, S5A). These data support the prediction that extensive mitochondrial reticulation sustains energetic balance in the shrew striated muscle.

Mitochondrial networks are extensive in *C. parva* striated muscle, with 75 ± 3.9% (skeletal muscle) and 67 ± 4.9% (cardiac) of the total mitochondrial volume participating in some form of matrix connectivity (Fig. 2N). In general, these mitochondrial interactions occupy small regions of the cell, while a few, large networks can occupy up to 19.3% of skeletal muscle and 13.6% of cardiac cell volume (Fig. 2P). This distribution suggests that these cells only require a few, large mitochondrial networks to distribute potential energy at the scale of the whole cell (Movie S4).

Mitochondrial nanotunnels allow distant mitochondria to maintain connectivity and are integral to the formation of the reticulum (25,27). We isolated nanotunnels from our mitochondrial segmentation and assessed the frequency and dimensions of these structures in skeletal muscle and heart. Mitochondrial nanotunnels are more common in cardiac muscle (16 ± 2.3 nanotunnels per 100 mitochondria) than skeletal muscle (5 ± 0.6 nanotunnels per 100 mitochondria) – and this high-frequency relative to human skeletal muscle (26) and mouse cardiomyocytes (24) is consistent with the large mitochondrial networks observed in *C. parva* (Fig. 3I). These nanotunnels have a diameter that is within range of the canonical < 200 nm diameter observed in other organisms (Fig. 3J) (28) and are morphologically distinct from the larger, non-cylindrical mitochondria that predominate in the I-band region of the skeletal muscle (Movie S3C - D). In *C. parva*, cardiac nanotunnels are shorter (mean = 354 ± 13.6 nm, max = 1.7 μm) than skeletal muscle nanotunnels (mean = 591 ± 40.8 nm, max = 1.6 μm) – which is likely due to the greater proportion of FPM in the heart resulting in shorter nanotunneling distances between mitochondria in intermyofibrillar (IMF) and paravascular regions (Fig. 3A-D) (19). When compared to larger organisms, these nanotunnels are of comparable length which suggests that there are structural or diffusive constraints limiting maximum nanotunnel length across eukaryotes (24,26,28).

### Liver mitochondria exhibit extensive interactions with the endoplasmic reticulum

We demonstrate that *C. parva* liver mitochondria do not form mitochondrial reticula like those found in the striated muscle of a variety of mammalian species. But *C. parva* liver mitochondrial structure is distinct as a result of an extremely high mitochondrial volume density (Fig. 2M), extensive interactions between mitochondria and the endoplasmic reticulum (ER) (Fig. 1C, 4) and specialized plasma membrane interactions (Fig. 4). We segmented the liver ER (Fig. S2) and quantified the percentage of outer mitochondrial membrane surface area within 50 nm of this organelle – the minimum distance required to detect a significant interaction between these surfaces (Fig. 4A) and is consistent with mitochondria-ER interactions in mouse hepatocytes (29). Based on these criteria, we show that 99.9% of all liver mitochondria have some ER interaction – a level that vastly exceeds the 20% described in HeLa cells and is similar to the frequency of interactions observed in mouse hepatocytes (29,30). This extensive association between mitochondria and the ER (mitochondrial-associated membrane; MAM) suggests that the ER could be the source of the high hepatic basal metabolic demand, driving the remarkably high concentration of mitochondria in *C. parva* liver (Fig. 2M). MAM has been identified as a site of mitochondrial phospholipid biosynthesis and Ca^+2^ regulation (30,31). But it is alternatively possible that the extensive MAM observed in *C. parva* acts to maximize ATP transport – or other metabolite – to the ER, fueling energetically expensive processes such as protein turnover and systemic lipid homeostasis (29,32). We demonstrate a positive correlation (y = 1.56x + 0.87, R^2^ = 0.64, p < 0.0001) between mitochondrial volume and the surface area of MAM (Fig. 4B). Thus, it appears that *C. parva* liver mitochondrial content can be adjusted to meet local energy or synthetic requirements – a phenomenon that is not observed in striated muscle.

**Figure 4.**
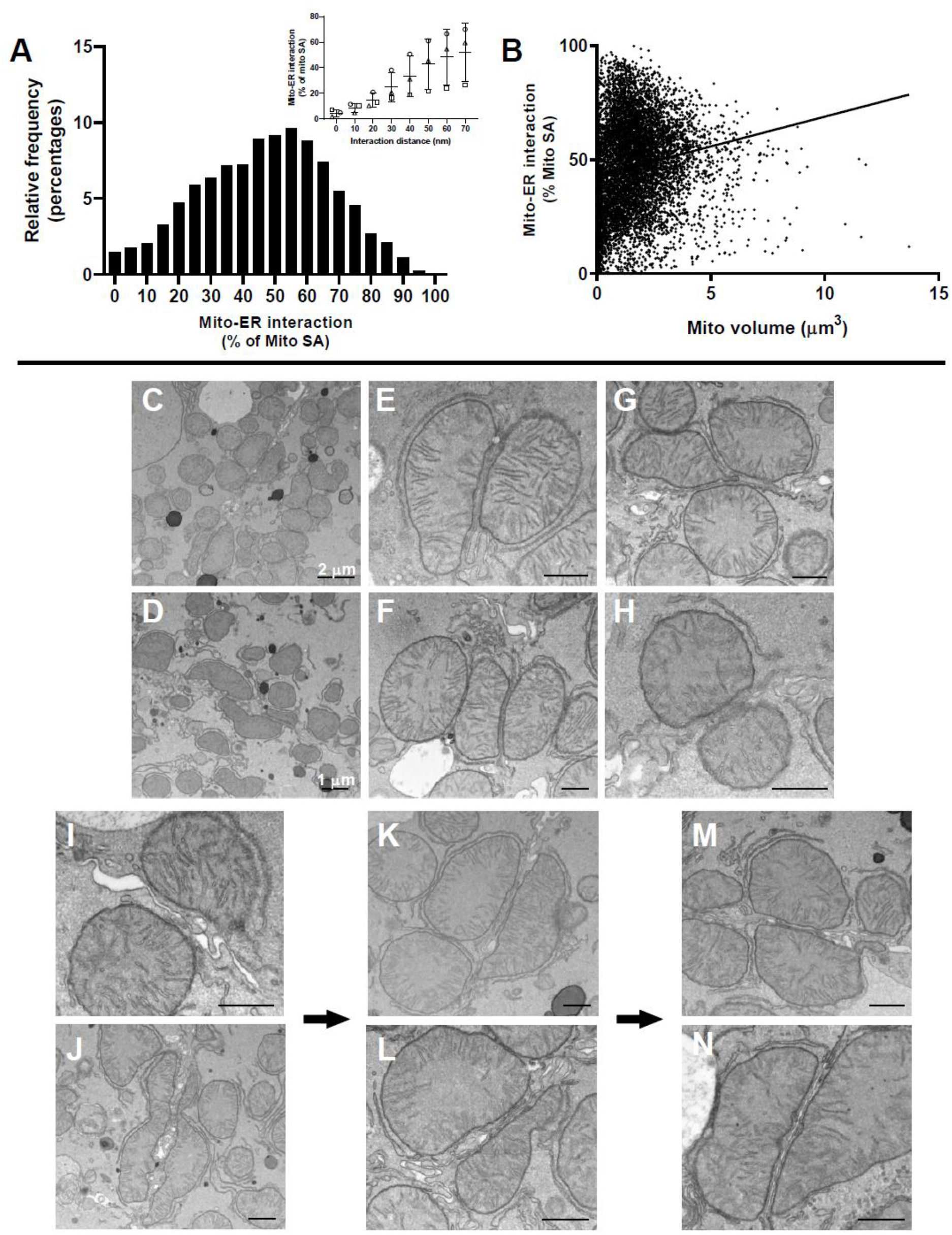
Liver mitochondria exhibit extensive inter-organelle interactions. **A**) Frequency distribution of individual mitochondrial surface area (percent of total) within 50 nm of the ER – the minimum distance required to detect a significant interaction between these organelles (subset figure, data are mean ± SEM; asterisks indicate significant difference from 0 nm, Dunnett’s multiple comparison, p < 0.05). **B**) Individual mitochondrial volume is positively correlated with mitochondrial associated-ER surface area (simple linear regression, y = 1.56x+0.87, R^2^ = 0.64, p < 0.05; *n* = 8941 mitochondria, three animals). **C-H**) Liver mitochondria exhibit membrane contacts across the plasma membrane (PM; paracellular membrane contact, PcMC). PcMC formation involves 1) mitochondria aligning at the PM and deformation of the outer mitochondrial membrane (**I-J**), 2) recruitment of ER between mitochondrial membranes and the PM (**K-L**) and 3) formation of the PcMC (**M-N**). Data are TEM images, scale bar: 500 μm unless otherwise indicated.

In addition to extensive MAM, we demonstrate that liver mitochondria in *C. parva* interact between cells across the plasma membrane (paracellular mitochondrial contact; PcMC; Fig. 4C-H; Movie S6). We identified three putative stages of the formation of PcMCs: 1) alignment of mitochondria at the plasma membrane and deformation of the OMM (Fig. 4I - J) 2) greater association between the mitochondria and PM that involves the recruitment of ER between the OMM and plasma membrane (Fig. 4K - L) and 3) formation of the PcMC (Fig. 4C - H, M - N). PcMCs are not loose associations as these structures remain intact following poor sample fixation and loss of other paracellular contacts (Fig. S3). The function of PcMCs remains unknown but similar structures have been described in human liver and bovine mammary glands and are proposed to involve desmosomes as part of their organization (33,34). The presence of at least four phospholipid bilayers (two sets of outer mitochondrial and plasma membranes) at these contacts makes it unlikely that they are electrically conductive as predicted to occur across muscle IMJs. But specialized channels may be present in these contact zones that are not directly observed in EM data. Alternatively, PcMCs may facilitate diffusive communication between mitochondria via small, soluble molecules such as ROS or Ca^+2^ (35) – allowing for direct mitochondrial regulation beyond the confines of a single hepatocyte.

### Kidney mitochondrial localization and morphology is cell-type specific

Our data from *C. parva* kidney reveals greater heterogeneity and distribution of cell-types when compared to striated muscle and liver. These cells maintain considerable variation in mitochondrial morphology and connectivity that likely reflects the localization of function along the nephron. Ion transport is a major driver of kidney metabolism – an activity that primarily occurs in the cortex and outer stripe of the medulla through the basolateral membrane localized Na^+^/K^+^-ATPase (36). Utilizing structural markers, we identified cells belonging to the proximal and distal convoluted tubule (PCT, DCT), the thick ascending limb (TAL), and collecting duct (CD) (Movie S7). In renal segments with high basolateral membrane Na^+^/K^+^-ATPase activity – such as TAL and DCT – large elongated mitochondria were found to be highly co-localized with the invaginated basolateral membrane. Similar renal mitochondrial localization has been observed in other mammals (37). In contrast, regions with low Na^+^/K^+^-ATPase activity, such as PCT and CD have more spherical mitochondria distributed across the cytoplasm. Elongated mitochondria in the TAL and DCT exhibit some mitochondrial connectivity through IMJs (Movie S8). But unlike striated muscle, these interactions do not result in cell-scale mitochondrial networking.

Similar to our observations in the liver, we note the presence of MAM in the kidney PCT (Movie S9). But due to a limited sampling of each cell-type, we are unable to quantify the extent of these interactions. Given the structural similarity of MAM between these tissues it is reasonable to predict that ER-associated protein turnover occurs in the kidney and is thus an important contributor to basal metabolic demand – likely at levels lower than those required for ion transport (14). Kidney PCT mitochondria also exhibit many instances of putatively failed mitochondrial fission (Movie S9). These events were identified based on a continuous OMM that runs through the interior of the organelle and a slight invagination of the OMM at the cytosolic interface. Notably, failed fission events were not observed in the liver and cardiac muscle in *C. parva* despite these tissues maintaining a similar mitochondrial volume density.

### Mitochondrial oxidative phosphorylation protein programming in the liver and kidney approaches levels observed in the heart of *Cryptotis parva*

The extreme mitochondrial volume density exhibited in the liver and kidney likely sustains basal metabolic demand in *C. parva*. But high mitochondrial content cannot support metabolism if it is not matched with sufficient protein programming to drive ATP synthesis. We compared the proteomes of heart, skeletal muscle, brain, liver and kidney and focused our attention on mitochondrial pathways, as shifts in protein abundance and composition likely allow *C. parva* to maintain energetic balance under extreme demand and enable tissue-specific mitochondrial processes beyond ATP synthesis (Fig. 5; *SI* appendix, data S1). In addition, we sequenced and annotated the *C. parva* genome to improve the accuracy of the proteomics database used in this comparison and to provide a genetic reference for this emerging research model (GenBank accession no.: To be submitted). The abundance of mitochondrial TCA cycle, β-oxidation, and ETS enzymes is generally low compared to the heart across sampled tissues (Fig. 5C – E). But this effect is smallest in the liver and kidney which is consistent with our spectrophotometric quantification of cytochrome *c* (Cyt *c*) and cytochrome a (Cyt a; Fig. 5A - B) content. When combined with tissue-specific variation in mitochondrial volume density (Fig. 2M) it appears that oxidative phosphorylation capacity in the kidney and liver is only marginally lower than the heart. This putatively high ATP synthetic capacity in liver and kidney likely supports the high metabolic demand of these tissues in *C. parva* and is consistent with increasing Cyt a content in the liver – but not the heart – with decreasing organism size (11).

**Figure 5.**
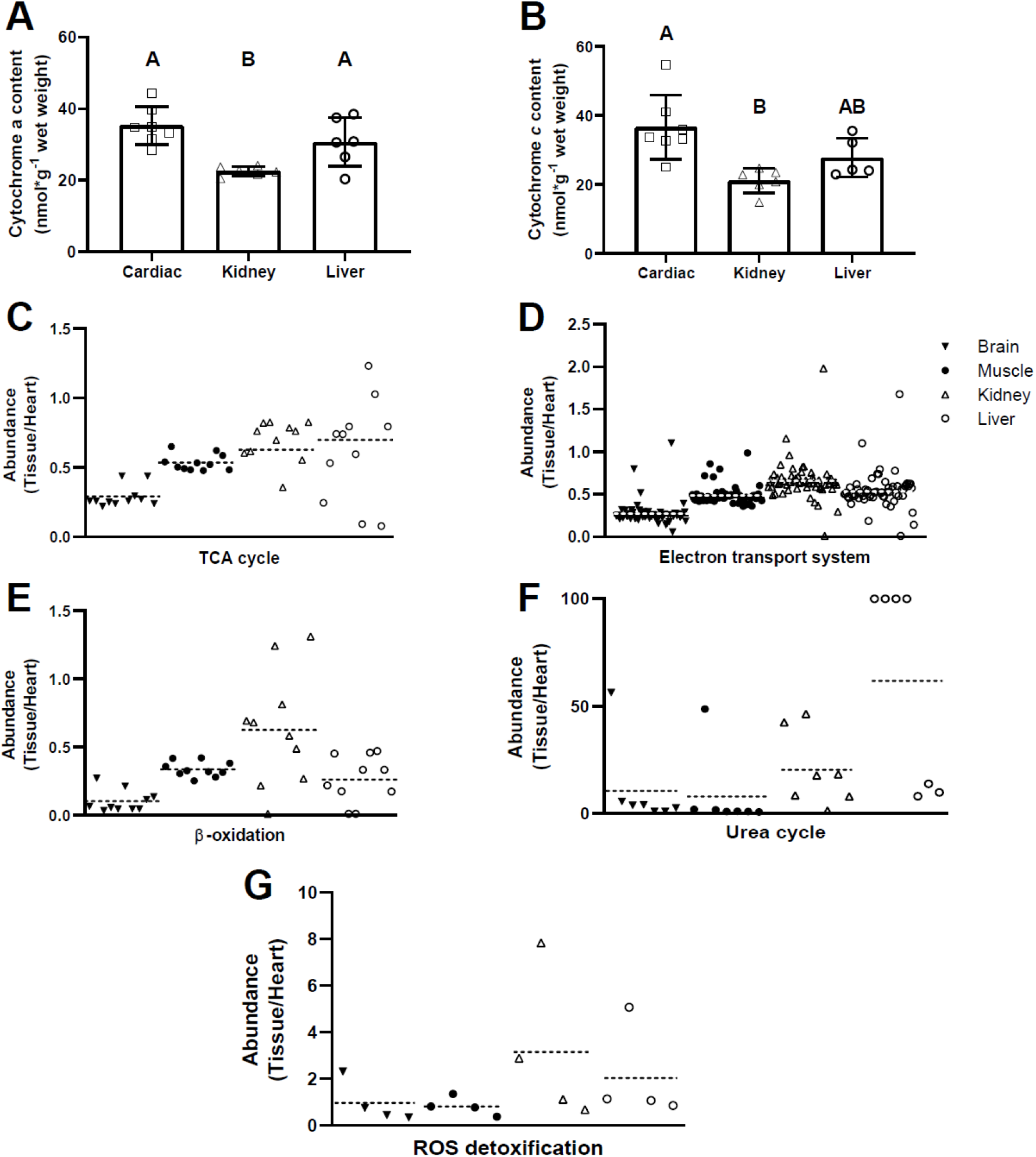
Liver and kidney mitochondrial protein programming approaches that of the heart in *Cryptotis parva*. **A**) Whole tissue cytochrome a content. **B**) Whole tissue cytochrome *c* content. Data are mean ± SEM and different letters indicate a significant difference between tissues (ANOVA, p < 0.05, *n* = 5-7 animals). **C-G**) Heart normalized protein abundance for biochemical pathways associated with mitochondria. Individual protein subunit abundance (datapoints) can be found in Data S1. Dashed lines indicate mean abundance value within a tissue. *n* = 4 animals.

Mitochondria are integral to mammalian urea cycling and ROS metabolism. Mitochondrial urea cycling occurs primarily in the liver although some steps of this process take place in the kidney (38) and this is supported by our data and a similar analysis in *Rattus norvegicus* (Fig. 5F) (39). Protein synthesis and degradation are energetically expensive and are thus major contributors to basal metabolism (4). Protein turnover has been proposed as a driver of the high basal metabolic rates in small organisms (40,41) and our findings – in conjunction with the extensive interactions between liver mitochondria and ER (Fig. 4) – support this prediction (42). But it should be noted that our relative protein abundance data only provide circumstantial evidence of high urea cycling capacity in *C. parva* relative to larger organisms. It remains unclear why protein turnover would be greater in smaller organisms, but one possibility is an increased production of free radicals in these species resulting in greater protein damage (43). This damage can be mitigated through mechanisms as simple as ROS detoxification. Mitochondrial ROS detoxification enzymes are most abundant in the kidney and liver (Fig. 5G) and this tissue-specific pattern is consistent with the tissue abundance of TCA cycle, ETS and β-oxidation enzyme content (Fig. 5D-E) – providing some evidence for high mitochondrial free radical generation and compensatory detoxification in *C. parva*.

## Discussion

In this study, we investigated mitochondrial content, structure and protein programming as components of the metabolic design of *C. parva*. The extreme volume density and localization of mitochondria in liver and kidney reveal solutions to the distribution of cellular energy conversion that are distinct from those observed in striated muscle. In striated muscle, the organization of mitochondrial reticulation conforms to patterns observed in other mammalian systems, with extensive networks – throughout the cytoplasm – that incorporate slightly enlarged nanotunnels and putative connectivity through IMJs. Cardiac mitochondrial networks are remarkably similar to those observed in the extensively studied mouse – with the exception that they maintain a greater incidence of nanotunnels. In contrast, skeletal muscle mitochondrial network structure differs when compared to larger mammals. For example, skeletal muscle mitochondria in the I-band region are not cylindrical, but instead form flattened sheets (Fig. 3A-B, Movie S3C-D) resulting in an increased surface area to volume ratio. This structure – combined with an individual IMF mitochondrial volume (0.37 ± 0.028 μm^3^, 2583 mitochondria, seven datasets, three animals) that is greater than that of the mouse (19) and dense cristae (Fig. S1, S4) – likely increases ATP synthetic capacity and the surface area for adenylate and phosphate transport adjacent to the myofibril. Skeletal muscle mitochondrial networks are also unique in their coupling of mitochondria through mitosynapses – primarily in the IMF region (Fig. 2I-L) – that increase surface contact between adjacent mitochondria. But the membrane and channel composition of mitosynapses and IMJs remains to be determined.

It is important to note that extensive mitochondrial reticulae – like those in striated muscle – are not observed in the liver and kidney, despite mitochondrial volume density in these latter tissues being similar to the heart (Fig. 2M). Mitochondrial content is generally matched to tissue metabolic demand (44), thus the high mitochondrial volume density in the liver and kidney indicates that the metabolic capacity in these tissues approach that of the heart under basal conditions. These data lend support to the proposed role of visceral organs – such as the liver and kidney – as major contributors to allometric scaling of basal metabolic rates (12–14,45). Given that mitochondrial volume density and thus energetic demand are similar among visceral organs it is intriguing that mitochondrial networks are absent in the liver and kidney. Our demonstration of consistent associations between mitochondria and other cytosolic structures in kidney and liver provides insight into an alternative structural solution to support high ATPase activity in these cells. In the mitochondrial rich cells of the kidney, mitochondria are embedded in the basolateral membrane of the epithelia where the Na^+^/K^+^-ATPase – a major energetic sink – is located. In the liver, mitochondria are stoichiometrically matched and tightly associated with the ER – an inter-organelle interaction that is putatively involved in many hepatic energy consuming processes (30,31). Thus, in these tissues, mitochondrial content and cellular energetic sinks are arguably matched – which provides a local balance between energy conversion and utilization and mitigates potential risks associated with electrically coupling mitochondria (24). This contrasts with striated muscle where the diffuse, high concentration of ATPase activity and the mechano-spatial requirements for contraction (i.e., end-to-end continuity) make it difficult to provide local energy supply. Consequently, mitochondria are positioned in areas that do not interfere with muscle mechanics (i.e., paranuclear, paravascular, and intramyofibrilar pools) (23) – which necessitates the formation of a mitochondrial reticulum to rapidly support the diffuse energetic demand of this tissue (17,24). The hummingbird – an avian model of extreme metabolism – potentially represents another extreme case of mitochondrial networking as 40% of flight muscle mitochondria are localized in the cell periphery (46). Thus, structural limitations – and not cellular energetic demand – appear to be the basis for the formation of cell-scale mitochondrial networks.

Cardiac mitochondrial volume density and maximum cardiac tissue respiration rate remain largely unchanged with decreasing mammal size, implying an optimization of cardiac peak sustained power and energy conversion across species (11,18). In addition, resting cardiac heart rate and associated metabolic rate are greater in small mammals – resulting in a loss of dynamic range for work and tissue metabolism as these organisms are forced to operate near their maximum metabolic capacity under basal conditions (18). This conclusion is supported by our data from *C. parva* as this species exhibits one of the highest mammalian basal metabolic rates and a remarkable heart rate, but limited alterations to cardiac mitochondrial content and structure relative to larger organisms (11,18,19). As a result, mammalian cardiac design appears to have approached an optimum that balances mitochondrial volume density and myofibril content in order to sustain muscle contraction at V_max_. In contrast, liver and kidney mitochondrial content increase with decreasing organism size suggesting that these tissues maintain greater metabolic scope despite large increases in basal metabolic rate (11).

Our understanding of the functional and evolutionary importance of mitochondrial morphology and networking is still in its infancy. Using data from the extreme case of *C. parva* – where the metabolic capacity of visceral organs approaches that of the heart – we propose that mitochondrial localization and structure are determined by the cytosolic distribution of cellular functions. This results in a local balance of energy conversion and utilization (i.e., liver and kidney) or a mitochondrial network to distribute energy from constrained sites (i.e., striated muscle). The signaling and regulatory aspects that control these distinct energy distribution mechanisms remains to be elucidated but it appears that these metabolic designs are maintained across the mammalian allometric series.

## Materials and Methods

### Animals

All animal use was approved by the National Heart, Lung and Blood Institute Animal Care and Use Committee (Protocol: H-0322). Least shrews (*Cryptotis parva*; 32 −38 day old males; whole animal mass = 4.7 – 6.0 g) – from a captive breeding colony (Western University of Health Sciences) – were housed under a 12:12 L:D cycle at 22 ± 1 °C. Animals were fed *ad libitum* with free access to water (47). Prior to sampling, all animals were held for two hours in clean plastic containers without food.

### Sample preparation for FIB-SEM

At the time of sampling, shrews were given an IP injection of heparin (2 μL, 1000 unit/mL stock) and were allowed to rest for 10 min. Animals were euthanized by exposure to isoflurane (32%). In order to fix the skeletal muscle, both hindlimbs were skinned and the exposed skeletal muscle was bathed in a fixative solution (2% formaldehyde, 2% glutaraldehyde in 0.1 M sodium phosphate buffer, pH 7.2) with the animal resting ventral side up. The heart, liver and kidney were fixed by first opening the chest cavity and cutting the right atrium. The heart was then perfused with 5 mL of relaxing solution (80 mM potassium acetate, 10 mM potassium phosphate, 5 mM EGTA, pH 7.2) using a peristaltic pump (flow rate = 3 mL/min) through the apex of the left ventricle, followed by a 1 h perfusion with fixative solution. During the perfusion fixation, exposed organs were bathed in fresh fixative solution that was exchanged every 15 min. Following the initial fixation, tissues were excised and placed in fixative solution overnight. Tissues were cut into 700 μm slices with a vibratome (VF-200, Precisionary Instruments, Boston, MA) and placed in fixative solution overnight.

Tissue slices were post-fixed and stained *en bloc* according to Bleck et al. (19) with slight modifications. Sliced tissues were placed in a standard fixative solution (2% formaldehyde, 1% glutaraldehyde, 0.1 M sodium cacodylate, pH 7.2-7.4) for 1 h followed by five washes of sodium cacodylate buffer (0.1 M; 3 min each). Samples were then post-fixed in a reduced osmium solution (1.5% potassium ferrocyanide, 0.1 M cacodylate, 2% OsO_4_) for 1 h on ice, followed by five washes with ddH_2_O (always 3 min each). Washed samples were next immersed in fresh 1% thiocarbohydrazide solution for 20 min at room temperature followed by five washes with ddH_2_O. Samples were post-fixed with a 2% OsO_4_ solution for 30 min on ice and washed five times with ddH_2_O. The sample was next incubated in 1% uranyl acetate overnight at 4 °C, washed five times with ddH_2_O, and incubated with freshly prepared Walton’s lead aspartate (0.02 M lead nitrate, 0.03 M aspartic acid, pH 5.5) for 30 min at 60 °C and washed again with ddH_2_O. The sample was then dehydrated using an increasing ethanol series (20%, 50%, 70%, 90%, 95%, 100%, 100%) at 5 min per step followed by incubation in 50:50 Epon:ethanol for 4 h and 75:25 Epon:ethanol overnight at room temperature. Finally, samples were incubated with 100% Epon in three steps of one, one and four hours and mounted in Epon according to the imaging technique.

Samples that were imaged using TEM and FIB-SEM were embedded in silicone molds and on aluminum ZEISS SEM mounts respectively and polymerized at 60 °C for 48 h. Epon mounted TEM samples were sectioned on an ultra-microtome and placed on 200-mesh grids prior to imaging. FIB-SEM samples were faced with a trim tool 90 diamond knife (DiATOME, Switzerland) and sputter coated with gold at a thickness of 50 nm (EMSX sputter coater, Electron Microscopy Services, Hatfield, PA).

### TEM imaging

TEM images were obtained using a JEM-1200EX electron microscope (JEOL USA) (accelerating voltage 80 keV) with a bottom-mounted 6-megapixel digital camera AMT XR111 digital camera (Advanced Microscopy Techniques Corporation, Woburn, MA). Final images were adjusted to balance contrast and brightness.

### FIB-SEM imaging

FIB-SEM samples were imaged with a ZEISS Crossbeam 540 FIB-SEM (Carl Zeiss Microscopy GmbH, Jena, Germany). Milling was performed with a FIB operating at 30 kV and 2-2.5 nA beam current. SEM images were collected at a voltage of 1.5 kV with a 1-1.5 nA of beam current and the in-lens detector captured backscattered electrons. The run and image stack alignment were controlled with Atlas 5 software (Fibics Incorporated, Ontario, Canada). Voxel size was set at 10 x 10 x 10 nm.

### Segmentation of cells and mitochondria

Image segmentation was completed on a desktop PC (Thinkmate, Waltham MA) running Windows 10 with Intel Xeon Gold 6254 3.10 GHz processors, 2.0 TB RAM and an NVIDIA Titan V 12 GB VRAM video card. We used Dragonfly software (Ver. 4.1; Object Research Systems, Montreal QC) and the U-net convolutional neural network (CNN) method (48) to segment organelles and cells.

To segment cells, we used a minimum of six training images per FIB-SEM volume. These training set images were down sampled three-fold (pixel size = 30 x 30 nm) in the image plane (XY volume orientation) and cell membranes were segmented through intensity thresholding followed by manual cleanup. In the case of the kidney, we excluded the brush border from the cell segmentation. We used these training datasets to build volume-specific cell-segmentation U-net models. In all instances, training data for U-net models were augmented (horizontal and vertical flip, 180° max rotation, 10° max shear, 75-150 % scale, 0-2 brightness, and 0-0.10 Gaussian noise) and 20% of the training data were used for validation. We utilized categorical cross entropy for our loss function and Adadelta for our optimization algorithm. The model was then applied to the full FIB-SEM volume to segment out cell membranes, which were then manually cleaned. To segment individual cells, we filled the inner area of the segmented cell membranes and ran a connected component analysis. We excluded data from cell volumes that were less than 1000 μm^3^ to ensure that we were not under-sampling.

Accurate segmentation of 3D organelles can be problematic as some CNN methods rely on segmentation of 2D images (e.g., U-net) – resulting in poor segmentation in orientations other than the image plane. To overcome this limitation, mitochondrial U-net models were trained using raw images derived from FIB-SEM volumes in three orientations (i.e., XY, YZ, and XZ; minimum three images per orientation). Outer mitochondrial membranes (OMM) were segmented from these training data using intensity thresholding and manual cleanup. The mitochondrial matrix was segmented by filling the inner area of the outer mitochondrial membrane. These training data were used to build volume-specific U-net models as described previously. We applied U-net models to each FIB-SEM volume in the same orientations described for the training data. The outer mitochondrial membrane and matrix were extracted from each reoriented U-net model segmentation and we created separate ‘triunion’ OMM and matrix segmentations using the union function – this was followed by manual cleaning. In order to segment individual mitochondria, we used a watershed mapping approach. Seeds for the watershed transformation were generated by subtracting the triunion OMM segmentation from the triunion matrix segmentation followed by manual cleaning and a connected component analysis. The boundary of the watershed was set by dilating the cleaned triunion matrix segmentation to include the OMM. From this segmentation we extracted mitochondrial surface area and volume and calculated SA/V, sphericity and the mitochondrial complexity index according to Vincent et al. (26) – with higher index values indicating greater complexity.

### Intermitochondrial junction segmentation

We employed our triunion U-net segmentation approach to segment intermitochondrial junctions (IMJ) in the skeletal muscle and heart. Previously described skeletal muscle and heart triunion U-net segmentation models were modified to include training against electron-dense IMJs as a separate ROI (Fig. 2A-L). We combined the triunion OMM and matrix segmentations, subtracted this union from the triunion IMJ segmentation and manually cleaned the resulting object. In order to quantify the extent of putative mitochondrial connectivity, we combined this IMJ segmentation with the mitochondrial matrix triunion segmentation and dilated the resulting object to include the OMM. Individual IMJ-linked mitochondrial pools were then individually segmented using a connected component analysis.

### Nanontunnel segmentation

We determined the prevalence and geometry of mitochondrial nanotunnels in skeletal and cardiac muscle using our previously described mitochondrial segmentation. Approximately 200 randomly selected mitochondrial meshes (STL format) per cell were simplified using Laplacian smoothing and two iterations of quadric edge collapse decimation in MeshLab (49). Resulting meshes were screened to ensure that they were two-manifold, closed and had no self-intersecting faces. Folded faces were removed, and any resulting disconnected components or holes were closed. Non-manifold vertices were repaired by splitting and resulting meshes were converted to the OFF format.

Mesh polygon counts were reduced through a custom command line C++ program (To be submitted to GitHub upon publication) that incorporates the triangulated surface mesh approximation function found in the Computational Geometry Algorithms Library (CGAL, 5.0.3). Holes and disconnected components in the resulting meshes were repaired through five iterations of removing disconnected components, closing holes, and repairing self-intersections. Each iteration of this repair was followed by removing faces of non-manifold edges. The final mesh was repaired through five iterations of repairing folded faces and splitting to repair non-manifold vertices.

Nanotunnel isolation and geometry (length, diameter) determination were completed using a custom C++ program (To be submitted to GitHub upon publication). Nanotunnel segmentation employed the shape diameter function in CGAL with default parameters. Mitochondrial meshes were skeletonized using the CGAL function with slight modifications to default parameters (medially centered speed tradeoff was set to 10, and the max number of iterations was set to 20). Meshes were segmented in to sub-meshes by the shape diameter function in CGAL with default parameters. Sub-mesh length was determined by generating a distance map from junctions separating adjacent mesh segments and calculating the shortest path connecting vertices nearest to the junctions using the Boost library. The longest Dijkstra shortest path was considered to be the submesh segment length. Sub-mesh diameters were determined by generating a series of planes orthogonal to the internal skeleton. The intersection between the plane and the sub-mesh segment were fit by a shell used to determine the min, max and average diameter of the sub-mesh segment. Nanotunnels were confirmed by filtering sub-meshes with a max diameter less than 300 nm and were manually checked against raw FIB-SEM data.

### Endoplasmic reticulum segmentation and MAM analysis

We quantified interactions between liver mitochondria and the endoplasmic reticulum (ER) – or mitochondrial associated membrane (MAM) – by first segmenting the ER using the triunion U-net segmentation approach. Liver mitochondrial U-net models were trained to include the ER, and the resulting ER segmentation was cleaned manually. We generated a distance map emanating from the ER and extracted surfaces based on the intersection of the distance map and the adjacent mitochondria. We then used Eqn. 1 to calculate the MAM surface area (SA_MAM_).

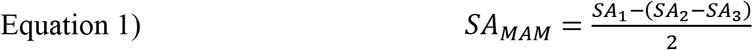

Where SA_1_ is mitochondrial SA, SA_2_ is SA of the mitochondrion excluding the volume of the intersecting ER distance map, and SA_3_ is SA of the region of the ER distance map that penetrates the surface of the mitochondrion. The MAM interaction distance for analysis was set at 50 nm (Fig. 4A).

### Sample preparation for genomics, proteomics and cytochrome quantification

Shrews were prepared for cardiac perfusion as described previously with the exception that hindlimbs were not skinned. Animals were perfused with 15 mL of ice-cold sodium phosphate buffer (0.1 M, pH 7.0) and the heart, brain, skeletal muscle, liver and kidney were excised. These tissues were dried of excess water on filter paper, weighed, and divided into smaller pieces for genomics, proteomics and cytochrome quantification. Tissue pieces were then snap frozen in liquid N2 and stored at −80 °C until later use.

### Genomic sequencing, alignment and annotation

We used a hybrid long and short-read genome sequencing approach to generate a genomic database for *C. parva*. High-molecular-weight genomic DNA was prepared from the kidney of one male shrew using a Genomic-tip 20/G (QIAGEN, Germantown, USA) according to the manufacturer’s protocol. The final genomic DNA preparation was suspended in TE buffer (0.1 mM EDTA).

### Long-read genome sequencing

The long-read sequencing library was constructed from 3 μg of genomic DNA using a Genomic DNA Ligation Sequencing Kit (Oxford Nanopore Technologies; New York, USA) following the manufacturer’s protocol. The ligated DNA library was loaded into a Flow Cell on MinION (Oxford Nanopore Technologies; New York, USA) for sequencing. Base sequence of the read was collected using MinKNOW software and processed into a read in the .fast5 format. The reads were transformed into basecalls and these results were written into FASTQ files, with a default of 4000 reads per FASTQ file.

### Short-read genome sequencing

Short-read sequencing libraries were constructed from 50 ng of genomic DNA using a Nextera XT DNA Library Preparation Kit (Illumina; San Diego, USA) following manufacturer’s protocol. The fragment size of DNAseq libraries was verified using the Agilent 2100 Bioanalyzer (Agilent; Santa Clara, USA) and concentrations were determined using a Qubit instrument (ThermoFisher Scientific; Waltham, USA). Libraries were checked for quality by sequencing on a MiSeq system (Illumia; San Diego, USA). The libraries were loaded onto the Novaseq 6000 (Illumia; San Diego, USA) for 2×50 bp paired end read sequencing. The FASTQ files were generated using the bcl2fastq software for further analysis.

### Canu assembly Long-read polishing with short-read sequencing

Contigs and unitigs (high confidence contigs) were generated from the long and short-read FASTQ files using the Canu pipeline (ver. 1.9) (50–53). Assembled contigs were polished by Medaka (version 0.12.1) to generate a consensus FASTA. The resulting genome was comprised of 39488 contigs with N50 of 112295 bp. Additional statistics for the Contig FASTA (generated from Canu) and consensus FASTA generated after medaka polishing can found in *SI* Table S1.

### Shrew genome annotation

The draft *C. parva* genome was compared with the human cDNA using BLAST (version 2.8.1) (54). Each BLAST hit was translated using TransDecoder (retrieved from http://transdecoder.github.io, version 5.5.0). The longest open reading frame (ORF) was selected for each cDNA, which also removed redundant blast hits. From these steps, 8659 shrew proteins were identified that also showed high homology to the human proteins.

### Tissue-specific proteomics

Snap frozen tissues (heart, brain, skeletal muscle, liver and kidney; n = 4 per tissue) were individually transferred to a lysis buffer (7 M urea, 2 M thiourea, 50 mM triethylammonium bicarbonate [TEAB]) at a 1:6 ratio and homogenized with ceramic beads using two steps of 20 s at 6000 rpm, 4 °C (Precellys^®^ Cryolys Evolution; Bertin Technologies, France). Tissue lysates were further homogenized using a QIAshredder homogenizer (Qiagen, Germantown, MD, USA) to reduce viscosity and clarified by centrifugation (15000 *g* for 10 min at 10 °C). Protein content in the supernatant was estimated using a Bradford assay (Thermo Fisher Scientific, Waltham, MA, USA) with BSA as a standard. Approximately, 150 μg of each lysate was digested with trypsin and labelled with Tandem Mass Tag (TMT) 11-Plex reagent kit according to the manufacturer’s instructions (Thermo Fisher Scientific, Waltham, MA, USA). In order to normalize across multiple TMT kits, we generated a pooled sample (15 μg from each lysate) internal control that was labelled with TMT label 131C.

### Offline HPLC peptide fractionation and Mass spectrometry

High pH reversed-phase liquid chromatography was performed on an offline Agilent 1200 series HPLC (Agilent Technologies, Santa Clara, CA, USA). The combined labeled digests (1.65 mg) were concentrated and desalted using two 1 cc HLB columns (Waters, Milford, CT, USA). The digest was resuspended in 0.1 ml 10 mM TEAB with 2% (v/v) acetonitrile. Peptides were loaded onto an Xbridge C_18_ HPLC column (Waters, Milford, CT, USA); 2.1 mm inner diameter x 100 mm, 5 μm particle size), and profiled with a linear gradient of 5–35 % buffer B (90% acetonitrile, 10 mM TEAB) over 60 min, at a flowrate of 0.25 ml/min. The chromatographic performance was monitored by sampling the eluate with an internal diode array detector scanning between wavelengths 200 and 400 nm, focusing on absorbances of 214, 254 and 280 nm. Fractions were collected at 1 min intervals followed by fraction concatenation (55). Fifteen concatenated fractions were dried and resuspended in 0.01% formic acid, 2% acetonitrile. Approximately 500 ng of peptide mixture was loaded per liquid chromatography-mass spectrometry run.

### Mass Spectrometry

All fractions were analyzed on an Ultimate 3000-nLC coupled to an Orbitrap Fusion Lumos Tribrid instrument (Thermo Fisher Scientific, Waltham, MA, USA) equipped with a nano-electrospray source. Peptides were separated on an EASY-Spray C_18_ column (75 μm x 50 cm inner diameter, 2 μm particle size and 100 Å pore size, Thermo Fisher Scientific, Waltham, MA, USA). Peptide fractions were placed in an autosampler and separation was achieved by 125 min gradient from 3-24% buffer B (100% ACN and 0.1% formic acid) at a flow rate of 300 nL/min. An electrospray voltage of 1.9 kV was applied to the eluent via the EASY-Spray column electrode. The Lumos was operated in positive ion data-dependent mode, using Synchronous Precursor Selection (SPS-MS3) (56). Full scan MS1 was performed in the Orbitrap with a precursor selection range of 380–1,500 m/z at nominal resolution of 1.2 x 10^5^. The AGC target and maximum accumulation time settings were set to 4 x 10^5^ and 50 ms, respectively. MS^2^ was triggered by selecting the most intense precursor ions above an intensity threshold of 5 x 10^3^ for collision induced dissociation (CID)-MS^2^ fragmentation with an AGC target and maximum accumulation time settings of 2 x 10^4^ and 75 ms, respectively. Mass filtering was performed by the quadrupole with 0.7 m/z transmission window, followed by CID fragmentation in the linear ion trap with 35% normalized collision energy in rapid scan mode and parallelizable time option was selected. SPS was applied to co-select 10 fragment ions for HCD-MS^3^ analysis. SPS ions were all selected within the 400–1,200 m/z range and were set to preclude selection of the precursor ion and TMTC ion series (57). The AGC target and maximum accumulation time were set to 1 x 10^5^ and 150 ms (respectively) and parallelizable time option was selected. Co-selected precursors for SPS-MS^3^ underwent HCD fragmentation with 65% normalized collision energy and were analyzed in the Orbitrap with nominal resolution of 5 x 10^4^. The number of SPS-MS^3^ spectra acquired between full scans was restricted to a duty cycle of 3 s.

### Data processing

Raw data files were processed using Proteome Discoverer (v2.4, Thermo Fisher Scientific, Waltham, MA, USA), with Mascot (v2.6.2, Matrix Science, Boston, MA, USA) search node. Due to the incomplete annotation of the *C. parva* protein database all peak lists were searched in parallel against the protein databases: UniProtKB/Swiss-Prot Homo sapiens (20,304 sequences, released 2020_7), NCBI reference sequence *Sorex araneus* (23,270 sequences, release 2020_7) and *Cryptotis parva* protein database derived from hybrid genome sequencing data (8,176 sequences) each concatenated with reversed copies of all sequences. The following search parameters were set with fixed modification carbamidomethylation of Cys, TMT 11-plex modification of lysines and peptide N-terminus; variable modifications of methionine oxidation and deamidation of Asn, and Gln. For SPS-MS3 the precursor and fragment ion tolerances were set to 10 ppm and 0.5 Da, respectively. Up to two-missed tryptic cleavages were permitted. Percolator algorithm (v.3.02.1, University of Washington) was used to calculate the false discovery rate (FDR) of peptide spectrum matches (PSM), set to a q-value <0.05 (58–61).

TMT 11-plex quantification was also performed by Proteome Discoverer by calculating the sum of centroided ions within 20 ppm window around the expected m/z for each of the 11 TMT reporter ions. All peptides were normalized and scaled to internal standard channels to reduce multibatch variability (62). Protein ratios were calculated separately, using the median pairwise ratio quantification for unique only (protein dependent isoforms) or shared peptides (protein group independent isoforms) at the MS^3^ level. The p-values were calculated for the reported quantitative ratios using analysis of variance (ANOVA) statistical testing.

### Pathway analysis for proteomics

Tissue proteome comparisons were limited to enzyme pathways associated with the mitochondrion (i.e., β-oxidation, TCA cycle, glycolysis, electron transport system, reactive O_2_ detoxification, and the urea cycle). We used human KEGG reference pathways (63) to identify proteins that belonged to each mitochondrial pathway of interest. We then compared the mean abundance ratio (each tissue relative to the heart) of all proteins within a given pathway to estimate the protein programming of each tissue.

### Cytochrome a and c quantification

We quantified liver, heart and kidney cytochrome *a* and *c* content spectrophotometrically (Shimadzu 2700 UV-Vis; Shimadzu Corp., Kyoto, Japan) based on established protocols (64). Tissue pieces were thawed on ice and homogenized (VirTishear Homogenizer, VirTis) in sodium phosphate buffer (0.1 M, pH 7.0). Tissue homogenates were solubilized with 1% dodecyl-β-maltoside followed by gentle mixing and centrifugation to remove solid material. The supernatant was used to measure difference spectra between reduced and oxidized states. The reduced state was achieved by incubating the sample with potassium cyanide (1 mM) and ascorbate (10 mM). Cytochrome *a* and *c* contents (molar extinction coefficients = 12 and 20.8 mM^-1^) were determined from the 605 and 550 nm wavelengths, respectively.

### Statistics

ANOVA testing was used to compare mean morphological differences and cytochrome content among tissues. When variances among groups were equal (determined by Welch’s t-test), a post-hoc Tukey test was used and when variances were not equal, a Dunn’s multiple comparisons test was applied. A t-test was used to compare mean morphological differences in between skeletal muscle and cardiac muscle under conditions where data were normally distributed (determined by the Shapiro-Wilk test). If these data failed the test of normality, then a Mann-Whitney test was applied.

The minimum distance required for a significant interaction between the liver ER and outer-mitochondrial membrane was determined using an ANOVA followed by a Dunnett’s multiple comparison test with all interaction distances being compared to the 0 nm interaction distance. Linear regression was used to compare the relationship between liver mitochondrial volume and the percent of mitochondrial OMM surface area that interacts with the ER.

Data analysis was completed using GraphPad Prism (9.0.0) with α = 0.05.

## Data Availability

Genome data will be submitted to NCBI GenBANK at the time of publication. All source data are available from the corresponding author upon reasonable request.

## Acknowledgements

The authors thank Heba Mohammed and Eric Lindberg for their electron microscopy technical expertise. The authors thank Mark Knepper for assistance with interpreting kidney data.

## Author Contributions

Conceptualization: DJC and RSB

Methodology: DJC, GPM, AMA, KS, MP, YL and CKB

Visualization: DJC

Funding acquisition: NAD and RSB

Supervision: RSB

Writing – original draft: DJC and RSB

Writing – final draft: all authors

## Materials and Correspondence

Requests for materials and correspondence can be made to DJC.

## Supplementary Materials

**Supplemental Figure 1.**
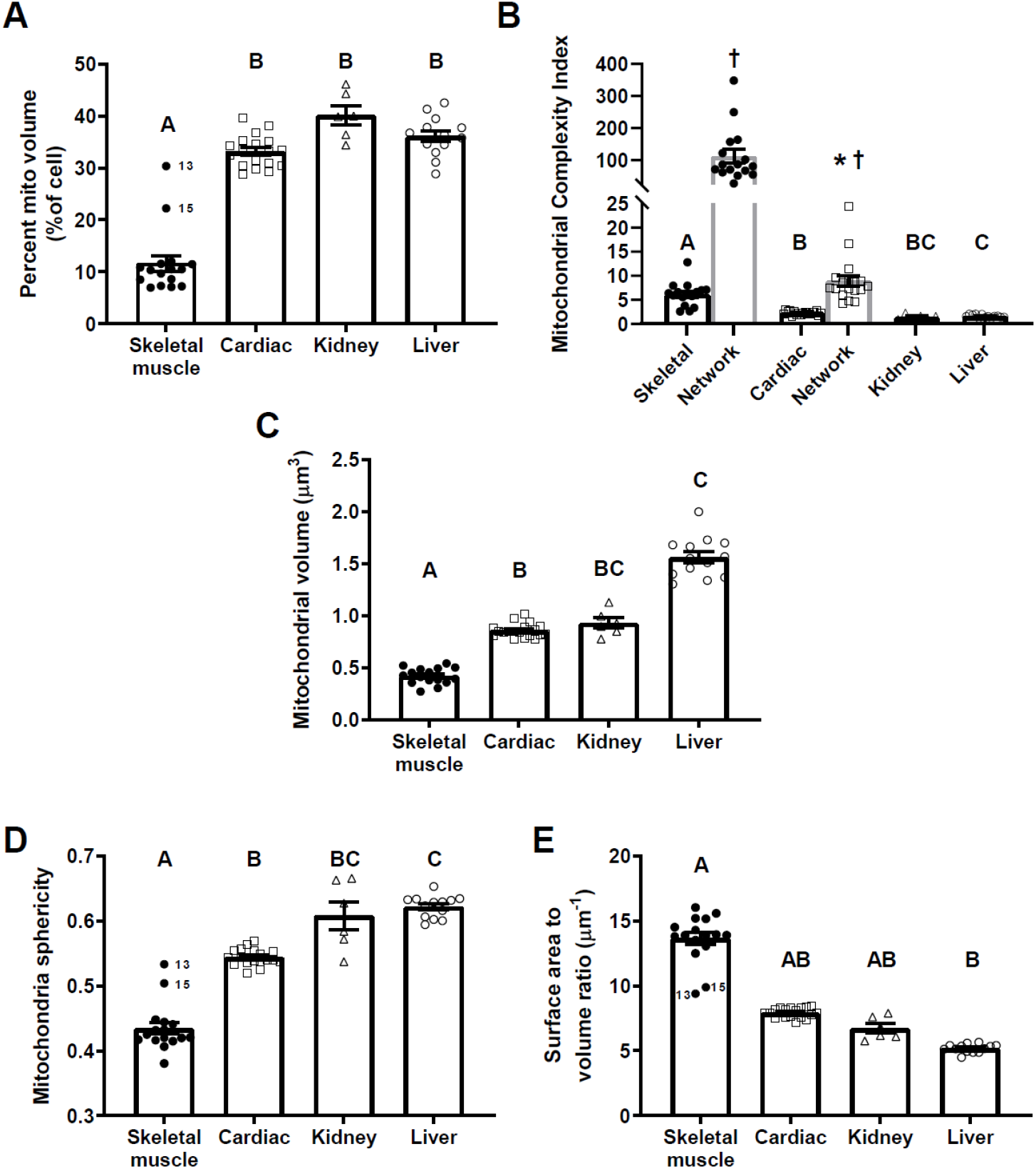
Morphological characteristics of mitochondria and mitochondrial networks in *Cryptotis parva*. (**A**) mitochondrial volume density. (**B**) Mitochondrial complexity index – an estimate of networking structure. (**C**) Individual mitochondrial volume. (**D**) Mitochondrial sphericity. (**E**) Mitochondrial surface area to volume ratio. Numbers are cell identifiers demonstrating the increase in estimated mitochondrial volume density associated with greater sampling of paravascular and paranuclear skeletal muscle mitochondria. Different letters indicate a significant difference among tissues (ANOVA, p < 0.05). Crosses indicate a significant difference between mitochondria and networked mitochondria within a muscle type (Tukey, p < 0.05) Asterisks indicate a significant difference between skeletal muscle and cardiac mitochondrial networks (Tukey, p < 0.05). Skeletal muscle, *n* = 19788 mitochondria, 16 datasets, three animals; cardiac muscle, *n* = 15893 mitochondria, 18 datasets, three animals; kidney, *n* = 16143, six datasets, two animals; liver, *n* = 6939, 13 datasets, three animals. Skeletal muscle networks, *n* = 242 networks, 16 datasets, three animals; cardiac muscle, *n* = 1218 networks, 18 datasets, three animals. Data are mean ± SEM.

**Supplemental Figure 2.**
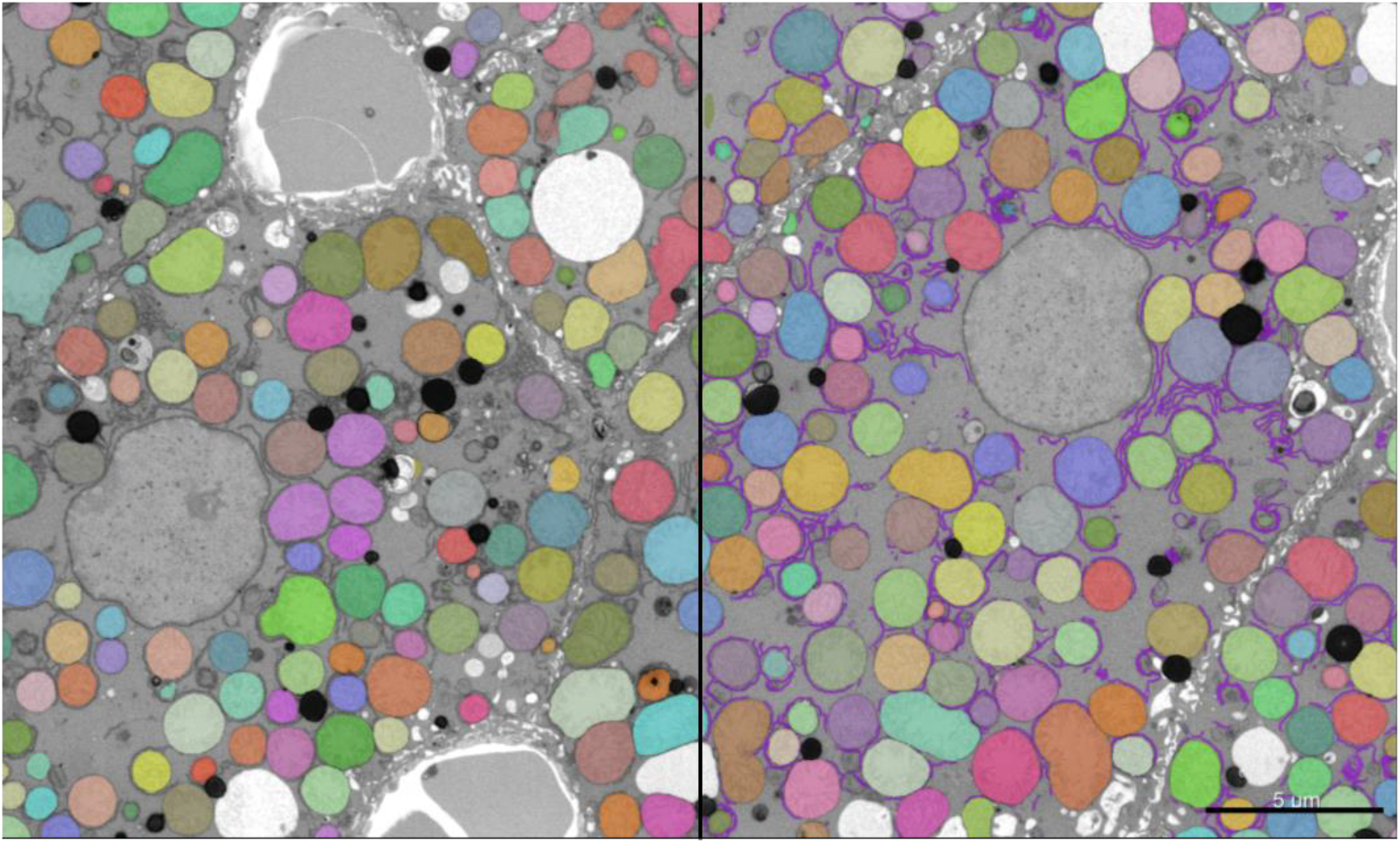
Representative endoplasmic reticulum (ER) segmentation. Data are raw FIB-SEM images (greyscale) with an overlay of mitochondrial (false color) and ER (magenta, image right-side) segmentation from *Cryptotis parva* liver. Distinct colors indicate individually segmented mitochondria. Scale bar: 5 μm.

**Supplemental Figure 3.**
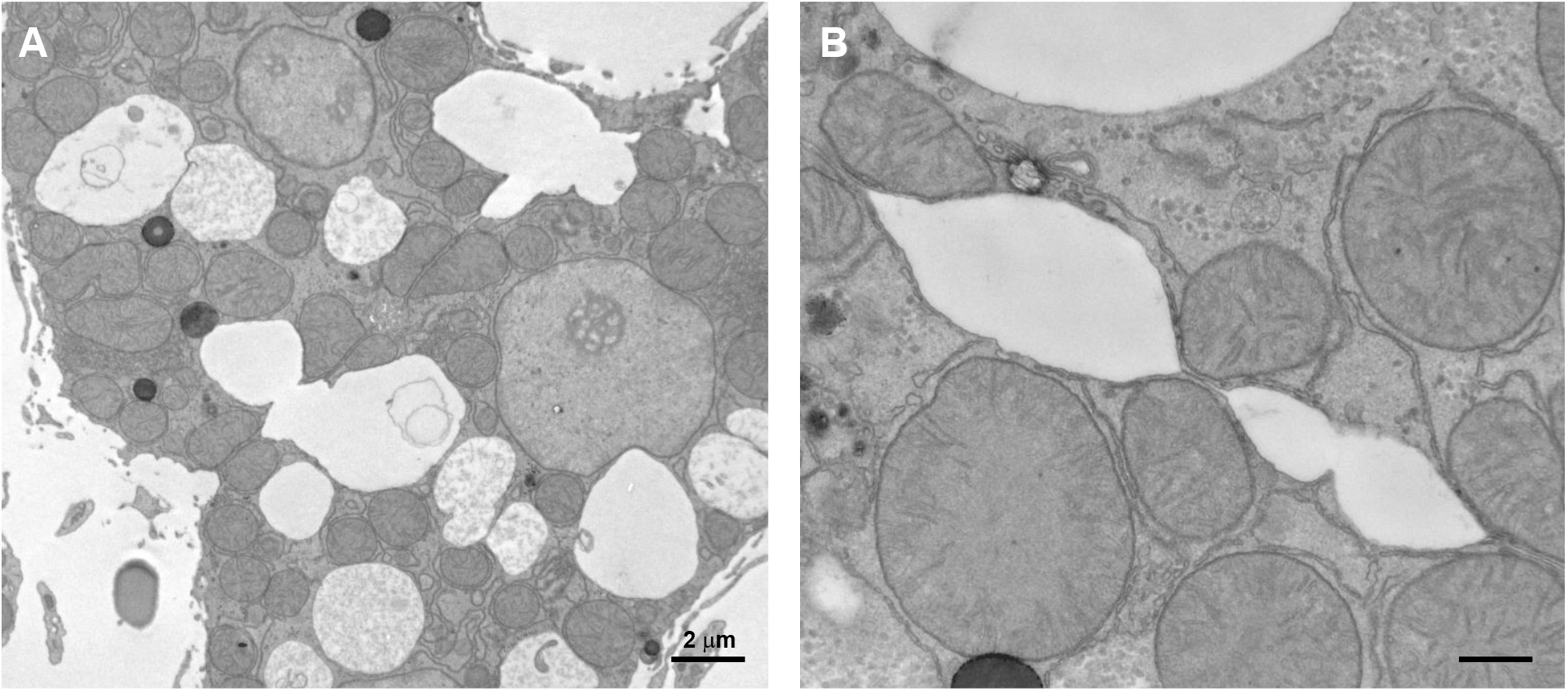
Liver paracellular mitochondrial contacts (PcMCs) are not loose associations. Poor sample fixation during cardiac perfusion results in delamination of adjacent plasma membranes. Despite poor fixation, PcMCs remain intact indicating that a structure holds these mitochondria in place. Data are raw TEM images. Scale bar: 500 nm unless otherwise indicated.

**Supplemental Figure 4.**
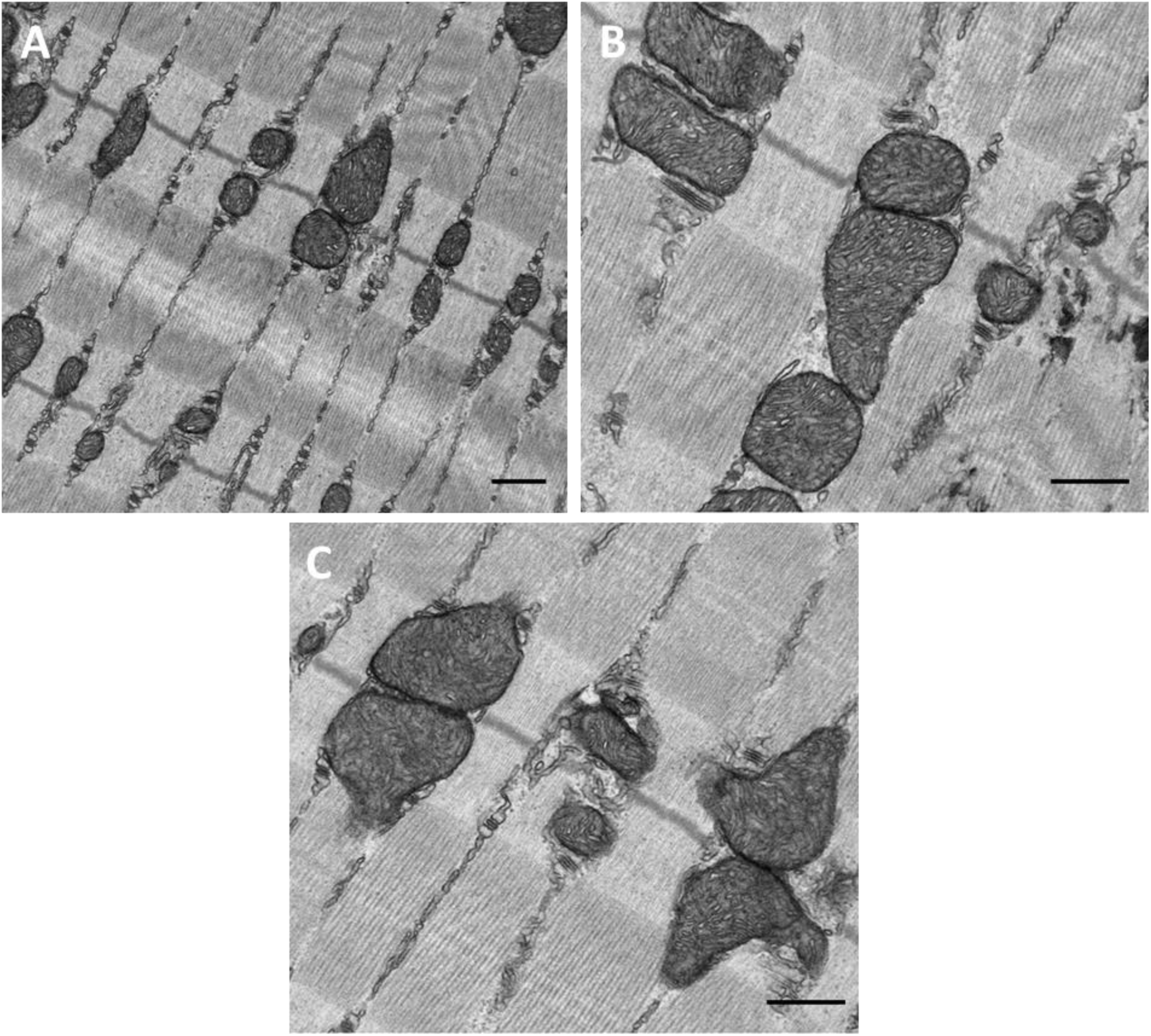
Skeletal muscle intermyofibrillar mitochondria have a high cristae surface area. Data are raw TEM images. Scale bar: 500 nm.

**Table S1:** Summary Statistics of Contig and Consensus FASTA files.

| Name | Contig FASTA | Consensus FASTA | Description |
| --- | --- | --- | --- |
| n | 41024 | 39488 | Total number of sequences |
| n:500 | 41024 | 39069 | Number of sequences at least 500 bp |
| L50 | 5247 | 5213 | Number of sequences at least the N50 size |
| min | 501 | 501 | The size of the smallest sequence |
| N80 | 51926 | 53332 | At least 80% of the assembly is in sequences of the N80 size or larger |
| N50 | 110332 | 112295 | At least half the assembly is in sequences of the N50 size or larger |
| N20 | 215680 | 218849 | At least 20% of the assembly is in sequences of the N20 size or larger |
| E-size | 163996 | 166513 | The sum of the square of the sequence sizes divided by the assembly size |
| max | 1612989 | 1631536 | The size of the largest sequence |
| sum | 2.18E+09 | 2.20E+09 | The sum of the sequence sizes |

## Supporting information captions

**Movie S1.**

Representative segmentation of mitochondria from four tissues in *Cryptotis parva*. Data are raw FIB-SEM image stacks (greyscale) with an overlay of mitochondrial segmentation (false color) from the heart (**A**), skeletal muscle (**B**), liver (**C**), and kidney (**D**). Distinct colors indicate individually segmented mitochondria. Scale bar: 5 μm.

**Movie S2.**

Representative segmentation of mitochondrial networks from cardiac (**A, B**) and skeletal muscle (**C, D**) in *Cryptotis parva*. Data are raw FIB-SEM image stacks with an overlay of individual mitochondrial segmentation (A, C) and mitochondrial network segmentation (B, D). Distinct colors indicate individually segmented mitochondria (A, C) or single mitochondrial networks (B, D). Scale bar: 10 μm.

**Movie S3.**

Representative cardiac (**A, B**) and skeletal muscle (**C, D**) intermitochondrial junction-linked mitochondrial networks in *Cryptotis parva*. Each volume represents a single IMJ-linked mitochondrial network. Distinct colors indicate individual mitochondria that comprise each network. Note the flattened-tube structure of the skeletal muscle I-band mitochondria (C, D).

**Movie S4.**

Cardiac (**A**) and skeletal muscle (**B**) mitochondrial network orientation in *Cryptotis parva*. Each volume represents a single EDZ-linked mitochondrial network (green) overlaid on a raw FIB-SEM volume (greyscale). Both volumes are oriented parallel to the plane of contraction. Note the parallel and perpendicular orientation of the cardiac and skeletal muscle mitochondrial networks respectively.

**Movie S5.**

Representative segmentation of mitochondrial intermitochondrial junctions (IMJ, magenta) from cardiac (**A**) and skeletal muscle (**B**) in *Cryptotis parva*. Segmented IMJs are overlaid on a raw FIB-SEM volume (greyscale). Both volumes are oriented perpendicular to the plane of contraction. Note the high IMJ content in cardiac fiber-parallel mitochondrial pools (A) and the high IMJ content in skeletal muscle paravascular mitochondrial pools (B). The central skeletal muscle cell (B) exhibits a high fiber-parallel mitochondrial content – commonly associated with a more ‘aerobic’ fiber type – which relies on IMJ linkage.

**Movie S6.**

*Cryptotis parva* liver mitochondria form paracellular mitochondrial contacts (white arrow). Data are representative raw FIB-SEM image stacks. Scale bar: 2 μm.

**Movie S7.**

*Cryptotis parva* kidney mitochondria from the cortex and outer stripe of the medulla exhibit a cell-type specific morphology and localization. The cortical proximal convoluted tubule (PCT) was identified based on the presence of the luminal brush border. The cortical distal convoluted tubule (DCT) identified based on the presence of a tightly packed cytosol with high mitochondrial volume density and nuclei that are close to one another. The outer-medulla thick ascending limb (TAL) was identified based on a high mitochondrial density and highly invaginated basolateral plasma membrane. The outer medullary collecting duct (CD) was identified based on the presence of a large central collecting vessel. Data are raw FIB-SEM image stacks. Scale bar: 10 μm.

**Movie S8.**

*Cryptotis parva* kidney mitochondria exhibit some connectivity through intermitochondrial junctions (identified by an increase in electron density between adjacent outer mitochondrial membranes, white arrow). Kidney mitochondrial connectivity is generally limited to cells that exhibit non-spheroid mitochondrial morphologies such as those in the thick ascending limb (TAL) but not the colleting duct (CD). Scale bar: 2 μm.

**Movie S9.**

*Cryptotis parva* proximal convoluted tubule kidney mitochondria interact with the endoplasmic reticulum (mitochondrial associated membrane) and exhibit failed fission events (white arrow). Data are representative raw FIB-SEM image stacks of cortical proximal convoluted tubule. Scale bar: 1 μm.

**Data S1. (separate file)**

Tandem mass tag proteomics results

